# Translating the Human CANTAB Touchscreen Based Tasks to Evaluate Learning and Memory in Mouse Models of Down Syndrome

**DOI:** 10.1101/2020.07.21.214106

**Authors:** Ashley Siegel, Diana W Bianchi, Faycal Guedj

## Abstract

Humans with Down syndrome (DS) exhibit hippocampal learning deficits in the Cambridge Neuropsychological Test Automated Battery (CANTAB). Here we translated the CANTAB Visual Discrimination (VD) and Extinction tasks to investigate hippocampal learning and cortical inhibitory control in the Dp(16)1/Yey, Ts65Dn and Ts1Cje mouse models of DS. No food or water restriction was used prior to testing. The number of days to reach 70% correct answers and percent of correct responses were analyzed. All Dp(16)1/Yey, Ts1Cje and WT mice reached Stage 5 of pre-training. No differences between genotypes were found in percent of correct responses. Five Ts65Dn and one WT animals reached Stage 5 and only one Ts65Dn mouse reached VD. Ts1Cje mice took longer (17.86±3.19 days) to move to VD vs. WT (11.44±1.96 days, *P*=0.09). There were no differences between Dp(16)1/Yey and WT mice. At VD, the average percent of correct answers was significantly lower in Dp(16)1/Yey (22.70±1.93%) and Ts1Cje (34.39±1.98%) compared to WT littermates (32.18±1.49% and 41.11±1.45%, respectively, *P*<0.05). In another set of experiments, we demonstrated that mild food restriction significantly reduced the time needed to complete pre-training in C57BL/6J mice compared to C67BL/6J mice that had *ad libitum* access to food and water. In conclusion, we were able to apply human cognitive tests to evaluate hippocampal learning and cortical inhibitory control in three mouse models of DS. These studies demonstrate significant cognitive differences between strains. Future experiments will evaluate whether food restriction and/or pre- and postnatal therapy decreases the time intervals to achieve training mile.

## Introduction

Microcephaly and associated intellectual disability are the hallmark of the brain phenotype in Down syndrome (DS) ^1^. In the last two decades, there has been a growing interest in developing therapeutic interventions to improve cognition and independent life skills in individuals with DS ^2^. Many pre-clinical trials, mostly conducted in the Ts65Dn mouse model, have shown significant improvement of brain and behavioral deficits in affected animals using a wide variety of compounds ^3–5^. Unfortunately, very few compounds were translated to human clinical trials and were largely unsuccessful ^6,7^. In two randomized double-blinded placebo trials, de la Torre *et al*. reported improvement in visual recognition memory, inhibitory control and adaptive behavior after treatment with epigallocatechin gallate (EGCG), a Dyrk1A inhibitor ^8,9^. Although promising, these studies must be extended to a multi-center trial with a larger number of participants and use more standardized testing batteries to build statistical power to evaluate outcomes.

The lack of success in translating preclinical findings to human clinical trials might be the result of multiple factors, including the choice of the best mouse models for pre-clinical studies, the timing of intervention (prenatal, early postnatal or adult s) and the lack of human translational cognitive outcome measures ^7^. The efficacy of therapeutic interventions is currently being evaluated using a wide range of behavioral tests (e.g. novel object recognition, Morris water maze and contextual fear conditioning for learning/memory), which do not have equivalents in human clinical studies.

When seeking to characterize the cognitive profile in humans with intellectual disabilities and speech delays, computerized tasks are widely used and are the most accurate methods of testing. Examples include the Cambridge Neuropsychological Test Automated Battery (CANTAB) and the National Institutes of Health (NIH) Toolbox Cognition Battery ^10–13^. These two testing batteries use non-verbal tasks that specifically target a particular brain region and offer the possibility to gradually increase the difficulty of each task. Important consequences of these characteristics include standardization and accessibility by a wide range of individuals and the ability to sensitively measure the severity of cognitive impairment. Additionally, the electronic “game-like” quality of these computerized tasks maintains interest and motivation more than paper tasks.

Recently, test batteries specifically designed for DS have been developed, including the Arizona Cognitive Test Battery (ACTB). Additionally, the NIH published their recommendations for certain outcome measures that are more appropriate to use in human clinical trials, among which the CANTAB and RBANS (Repeatable Battery for the Assessment of Neuropsychological Status) represent the main endpoint measures for learning and memory in humans with DS ^14^. Regarding tasks on the CANTAB, several studies demonstrate that individuals with SS exhibit significant deficits in Visual Discrimination and Paired Associates Learning (hippocampal-dependent learning/memory), Rule Reversal (frontal lobe-dependent learning flexibility) and CANTAB Simple Reaction Time or SRT (cerebellum-dependent processing speed) ^15–17^.

Here we used rodent touch screen behavioral paradigms to investigate hippocampal-dependent learning and frontal lobe inhibitory control in three different mouse models of DS (Dp(16)/1Yey, Ts65Dn and Ts1Cje). We hypothesized that trisomic animals would show learning/memory deficits compared to wild type littermate controls. We also expected, based on our previous comprehensive comparative analysis of phenotypes that there would be differences in performance among the three strains examined in this study ^18^.

The long-term goal of the current study is to establish human translational behavioral endpoints that can be used to analyze the effects of future prenatal and/or postnatal therapeutic interventions in pre-clinical trials.

## Materials and Methods

### Animals

All murine experiments were conducted in accordance with the National Institutes of Health Guide for Care and Use of Laboratory Animals and were approved by Tufts University Institutional Animal Care and Use Committee (IACUC) under protocol B2015-171. Animals were housed in cages with standard bedding and a nestlet square. The colonies were maintained on a 12-hour light/dark cycle, with lights on at 07:00 am. Two separate sets of experiments were conducted in this study.

For the first set of experiments, we used 4-5 month-old male mice from three different strains, including B6129S-Dp(16Lipi-Zfp295)1Yey/J [known as Dp(16)1/Yey] mice (Jax Stock number 013530), B6EiC3Sn.BLiA-Ts(17^16^)65Dn/DnJ [known as Ts65Dn] mice (Jax Stock number 005252) and B6.Cg-T(12;16)1Cje/CjeDnJ [known as Ts1Cje mice] (Jax Stock number 004838). For each strain, trisomic animals were compared to their WT littermate controls. The total number of animals used was as follows: 12 Dp(16)1/Yey versus 14 WT littermates, 12 Ts65Dn versus 12 WT littermates and 12 Ts1Cje versus 14 WT littermates. Genotyping was performed as previously described ^18^. No aversive stimuli (food or water restriction) were used in this first set of experiments in order to mimic the procedures used in human clinical trials.

For the second set of experiments, we used 10 C57BL/6J male mice (five weeks old) to evaluate the impact of mild food restriction on training length and motivation. Half of the mice underwent food restriction to achieve 85% of body weight, while the other half had access to food and water *ad libitum*.

### Apparatus

Mice were trained and completed behavioral tasks in a touchscreen-based apparatus consisting of a modular operant chamber within a sound and light attenuating chamber (Med Associates Inc. Fairfax, VT). Every operant chamber (Model ENV-307W-CT, length= 21.59 cm, width= 18.08 cm, height= 12.7 cm) is equipped with a touch screen on one end and a food/fluid receptacle equipped with a fluid dispenser, reward light, nose-poke detector, tone generator and house light on the other end. The touchscreen was overlaid with a black Plexiglas three-squared apertures (length=3 cm, height= 3cm) that allowed for a total of three visual stimuli to be displayed on the screen at one time (1 stimulus in each square aperture). The floor of the chambers consisted of thin aluminum bars spaced 0.5 cm apart. Chamber functioning, stimulus presentation and reward delivery were controlled through the K-Limbic software (Conclusive Solutions, Sawbridgeworth, UK).

### Pre-training

Mice completed one session every day for each Stage of pre-training in the same chamber. Each session lasted 30 min in all pre-training Stages except for Stage 5 which consisted of 60-minute sessions. Animals were individually advanced to subsequent Stages when they reached criterion regardless of the progress of other mice being tested.

#### Habituation

In order to habituate the animals to the reward (Chug chocolate milkshake), 200 uL of the milkshake was dispensed in a 35 × 10 mm Petri dish and transferred to the mouse home cage. Mice needed to consume the entire allotment of milkshake for two days in a row before they were advanced to Stage 1.

#### Stage 1

Mice were placed in the testing chambers for 30 minutes with 200 uL of chocolate milkshake in the liquid reward receptacle in order to accustom the mice to retrieving the reward. Completion of Stage 1 only occurred after all the milkshake was consumed within the allotted time frame two days in a row.

#### Stage 2

Stage 2 began the day following completion of Stage 1. During this Stage, mice were shown an image of a flower (CS+) in one of the three apertures while the other two apertures remained blank (inactive touch). The image remained on the screen indefinitely until a response was made. Responses to the screen within the empty apertures were not registered and led to no punishment, while a touch response to the CS+ ended the trial and resulted in the dispensing of 10 µL of milkshake into the receptacle, illumination of the receptacle light and a reward tone. Once the reward was collected, the inter-trial interval (ITI) of 10 seconds would begin, after which the flower image re-appeared on the screen and the trial was repeated. The position of the CS+ within one of the three apertures was randomly generated for each trial and did not follow any specific pattern. Criterion for advancing to the next Stage was 20 responses during the 30-minute session.

#### Stage 3

Stage 3 followed the same procedure as Stage 2 except that the CS+ remained on the screen for only 30 seconds rather than indefinitely.

#### Stage 4

Stage 4 commenced the trial with the illumination of the receptacle light which required mice to “nose-poke” into the receptacle to initiate a trial and presentation of the CS+. This time, mice had 20 seconds to make a response and touches to the black apertures were not registered. After the ITI, the receptacle light was once again illuminated signaling that the next trial could be initiated.

#### Stage 5

Stage 5 was the last Stage of pre-training and introduced a punishment when incorrect responses to the screen were made. Once again, trial initiation was required by nose-poking into the illuminated receptacle which lead to the appearance of the CS+. The image remained on the screen for 20 seconds and during that time, incorrect responses to the two blank apertures were registered and led to immediate removal of the CS+ and a 15-second timeout during which the house light was illuminated, and no trials could be initiated. Each timeout was followed by the ITI and a correction trial (CT) in which the CS+ appeared in the same aperture as the previous trial in order to prevent the mice from developing a side bias. CTs were repeated until a correct response was made. Criterion for advancing to the visual discrimination task was 70% correct responses during the 60-minute session.

### Visual Discrimination

After completing touchscreen pre-training, mice from the three strains moved to the Visual Discrimination (VD) task where they were taught to differentiate between a rewarded and an unrewarded visual stimulus; the images were a spider (CS+) and plane (CS-). The images were novel to the mice and were displayed simultaneously on the touchscreen in random positions for each trial. A session began with illumination of the receptacle light, indicating to the mouse to initiate the trial with a nose-poke into the receptacle. Once initiated, the CS+ and CS-appeared within two of the apertures and mice were expected to make a response. A nose-touch to the CS+ led to 10 µL milkshake reward, illumination of the receptacle and the reward tone. A nose-touch to either the CS- or the blank aperture led to a 15-second timeout during which time the house light was illuminated, no response could be made to the screen and no trial could be initiated. After the timeout, a 10 second ITI would commence followed by illumination of the receptacle light. This signaled that the next trial could be initiated. Incorrect trials were followed by a CT. Mice were trained at the VD task for 30 consecutive days. Each session lasted 60 minutes.

### Extinction

After completing VD, mice were placed in the Extinction phase for 10 consecutive days. A trial began in the same manner as VD and displayed the same CS+ and CS-after initiation. The difference was that responses made to the screen, whether correct or incorrect, resulted in no punishment or reward. Nose-touches to any part of the screen only led to the immediate removal of the images and illumination of the receptacle light with no milkshake reward even when the CS+ was touched. The ITI was once again 10 seconds.

### Statistical Analysis

The following dependent measures were recorded and analyzed at all testing Stages: the average percent of correct touches, the average number of days to reach criteria, the percent of mice reaching criteria every day, the percent of misses (in the Extinction sessions) and the average number of trials. For all measures, except the percent of mice reaching criteria every day, the Kolmogorov-Smirnov normality test and the Fisher test for variances comparison were first used to define the appropriate statistical test (parametric t-test or non-parametric Mann-Whitney test) for each dataset.

For the percent of mice reaching criteria each day, the area under the curve (AUC) was analyzed in trisomic versus WT littermates and compared using the unpaired t-test of AUC. Data were represented as mean ± SEM and statistical significance was reached with a *P*-values < 0.05. All statistical analysis and graphs were generated using GraphPad Prism 8.3.1 software package (GraphPad Software, San Diego, CA).

## Results

### I Comparison of performances in the Dp(16)1/Yey, T65Dn and Ts1Cje mouse models without food restriction

#### Pre-training

##### Early Pre-training (Stages 2-4)

All Dp(16)1/Yey mice and their WT littermates successfully completed early pre-training at some point throughout the experimental period (Figure 1–2 A, Table 1). There were no differences in the average number of days it took trisomic and WT animals to reach criteria at Stages 2, 3, and 4 (Figures 1–2 B). No difference was also noted in the average number of trials that were initiated by trisomic and WT mice at early pre-training Stages (Figures 1–2 C, Table 1).

For Ts65Dn mice, the average pecrentage of animals reaching criteria per day was significantly different between Ts65Dn trisomic and WT mice at all Stages of early pre-training, with more Ts65Dn animals advancing each day compared to their WT littermates (Figure 1–2 D, Table 1). There were no statistically significant differences in the average number of days it took both trisomic and WT mice to reach criteria at Stages 2 or 3, but only four WT and seven Ts65Dn trisomic animals ultimately advanced to Stage 4 (Figure 1–2 E, Table 1). However, the number of trials initiated at Stages 2 and 3 by Ts65Dn mice was significantly higher (Stage 2= 9.21 ± 0.70, Stage 3= 15.31 ± 2.06) compared to their littermate WT controls (Stage 2= 4.96 ± 0.57, *P*<0.001; Stage 3= 7.42 ± 1.33, *P*<0.01) (Figure 1F, Table 1). At Stage 4, only one WT achieved criteria and advanced to Stage 5, therefore we could not compare performances between cohorts.

**Table 1:**
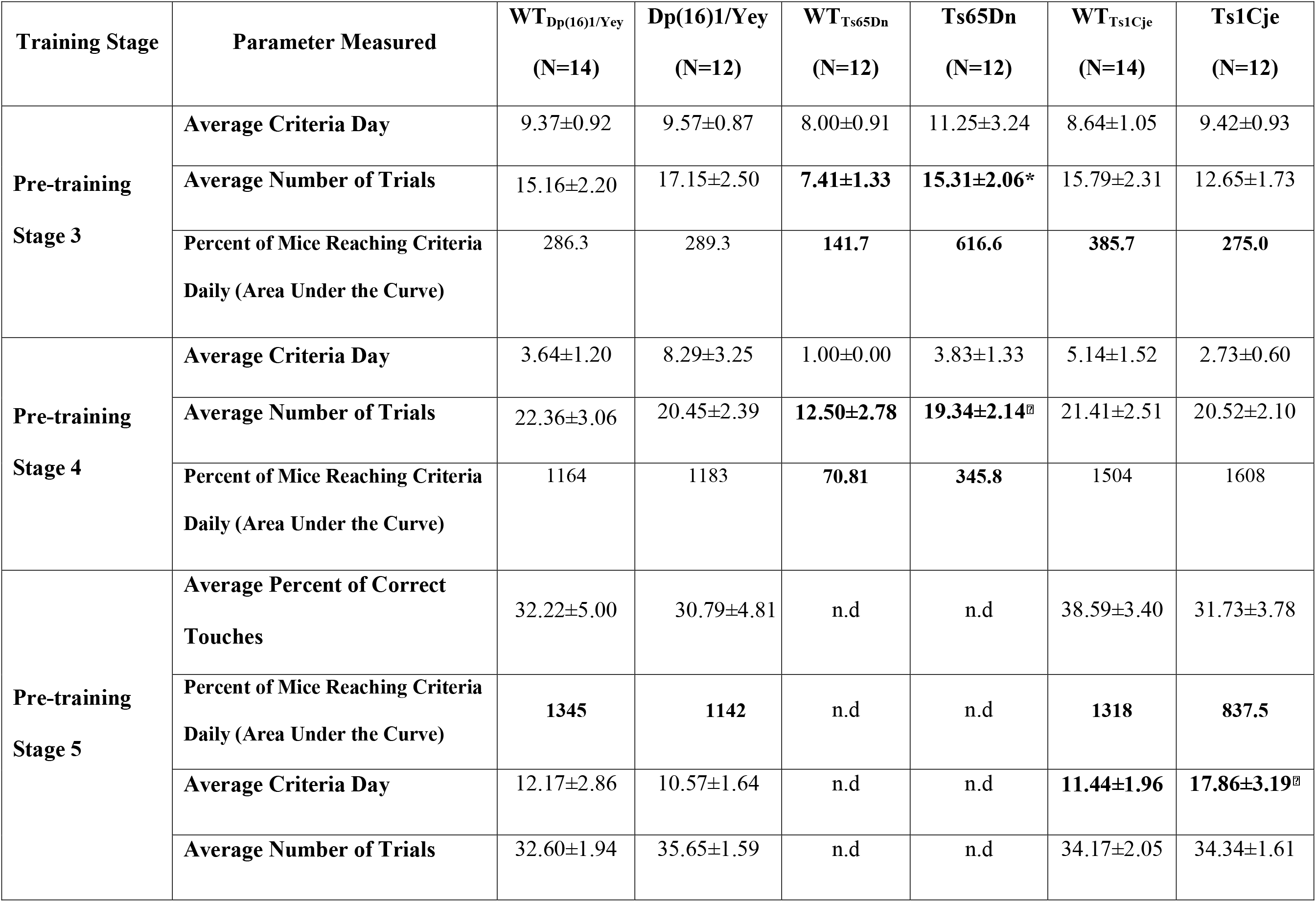
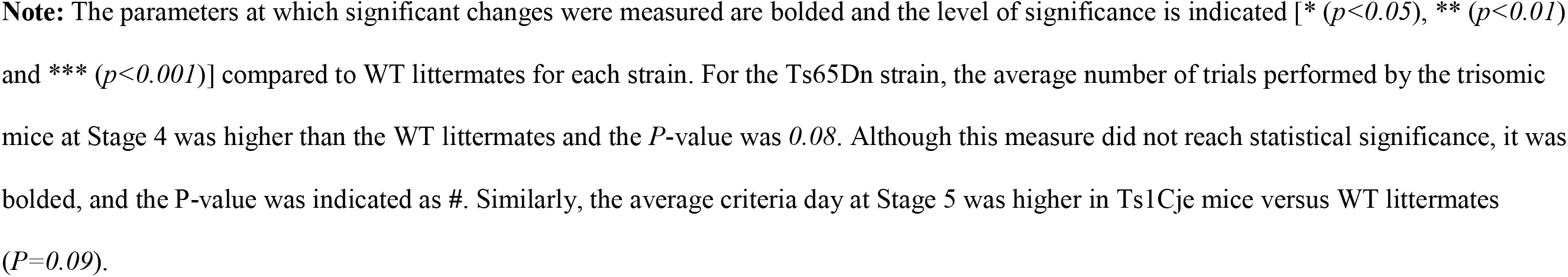
Pre-Training Results Summary in the Dp(16)1/Yey, Ts65Dn and Ts1Cje Mouse Models.

| **Training Stage** | **Parameter Measured** | **WT<sub>Dp(16)1/Yey</sub>**<br>**(N=14)** | **Dp(16)1/Yey**<br>**(N=12)** | **WT<sub>Ts65Dn</sub>**<br>**(N=12)** | **Ts65Dn**<br>**(N=12)** | **WT<sub>Ts1Cje</sub>**<br>**(N=14)** | **Ts1Cje**<br>**(N=12)** |
| --- | --- | --- | --- | --- | --- | --- | --- |
| **Pre-training**<br>**Stage 3** | **Average Criteria Day** | 9.37±0.92 | 9.57±0.87 | 8.00±0.91 | 11.25±3.24 | 8.64±1.05 | 9.42±0.93 |
|  | **Average Number of Trials** | 15.16±2.20 | 17.15±2.50 | **7.41±1.33** | **15.31±2.06\*** | 15.79±2.31 | 12.65±1.73 |
|  | **Percent of Mice Reaching Criteria**<br>**Daily (Area Under the Curve)** | 286.3 | 289.3 | **141.7** | **616.6** | **385.7** | **275.0** |
| **Pre-training**<br>**Stage 4** | **Average Criteria Day** | 3.64±1.20 | 8.29±3.25 | 1.00±0.00 | 3.83±1.33 | 5.14±1.52 | 2.73±0.60 |
|  | **Average Number of Trials** | 22.36±3.06 | 20.45±2.39 | **12.50±2.78** | **19.34±2.14<sup>§</sup>** | 21.41±2.51 | 20.52±2.10 |
|  | **Percent of Mice Reaching Criteria**<br>**Daily (Area Under the Curve)** | 1164 | 1183 | **70.81** | **345.8** | 1504 | 1608 |
| **Pre-training**<br>**Stage 5** | **Average Percent of Correct**<br>**Touches** | 32.22±5.00 | 30.79±4.81 | n.d | n.d | 38.59±3.40 | 31.73±3.78 |
|  | **Percent of Mice Reaching Criteria**<br>**Daily (Area Under the Curve)** | **1345** | **1142** | n.d | n.d | **1318** | **837.5** |
|  | **Average Criteria Day** | 12.17±2.86 | 10.57±1.64 | n.d | n.d | **11.44±1.96** | **17.86±3.19<sup>§</sup>** |
|  | **Average Number of Trials** | 32.60±1.94 | 35.65±1.59 | n.d | n.d | 34.17±2.05 | 34.34±1.61 |

**Note:** The parameters at which significant changes were measured are bolded and the level of significance is indicated [ $*$ ( $p < 0.05$ ), $**$ ( $p < 0.01$ ) and $***$ ( $p < 0.001$ )] compared to WT littermates for each strain. For the Ts65Dn strain, the average number of trials performed by the trisomic mice at Stage 4 was higher than the WT littermates and the $P$ -value was $0.08$ . Although this measure did not reach statistical significance, it was bolded, and the $P$ -value was indicated as $\#$ . Similarly, the average criteria day at Stage 5 was higher in Ts1Cje mice versus WT littermates ( $P = 0.09$ ).

**Figure 1:**
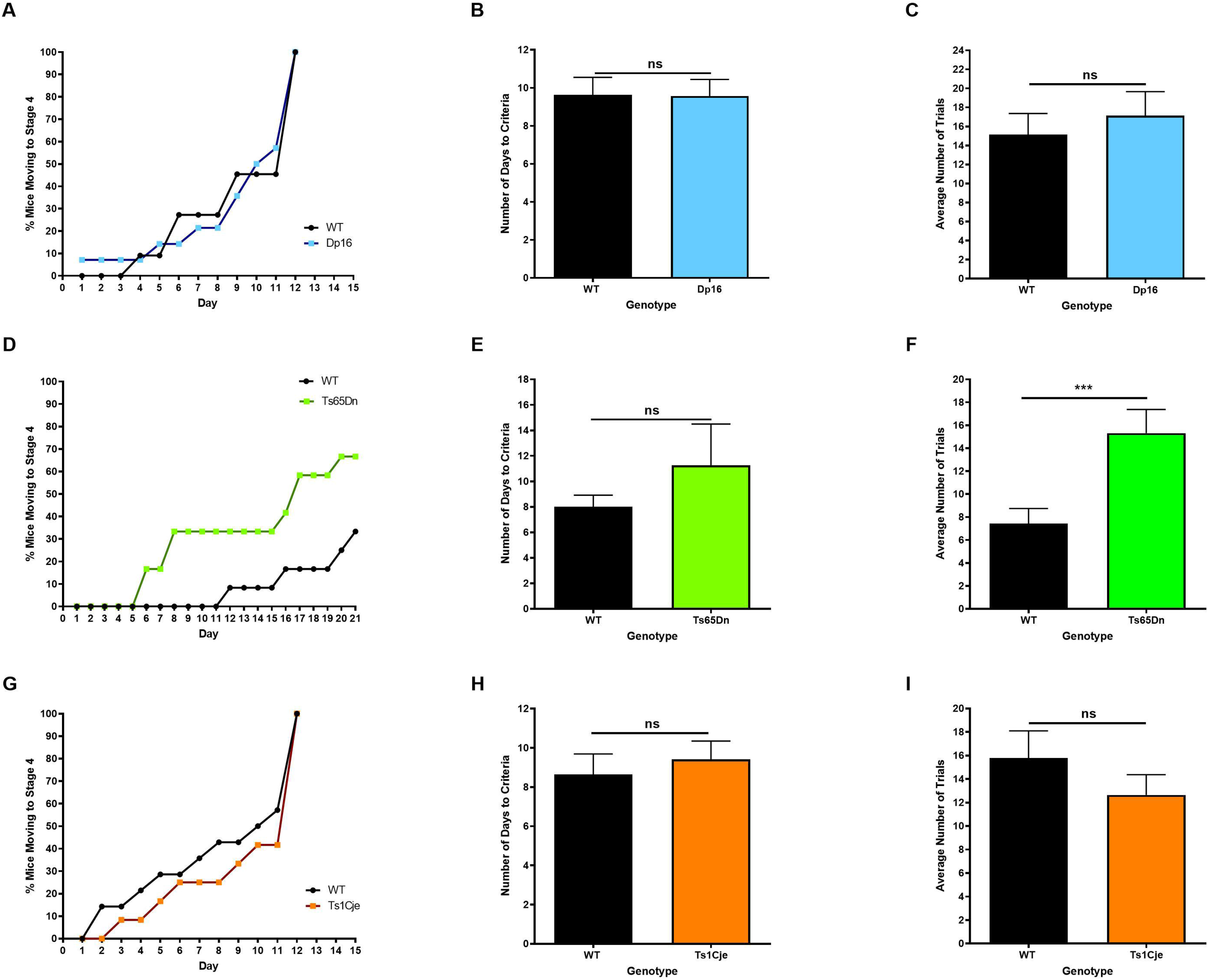
Dp(16)1/Yey, Ts65Dn and Ts1Cje Trisomic Mice Performances in Stage 3 of Pre- Training Compared to WT Littermates. **A-C:** The average percent of mice reaching criteria daily (A), average criteria day (B) and average number of trials (C) initiated by the Dp(16)1/Yey trisomic mice versus WT littermates. **D-F:** The average percent of mice reaching criteria daily (D), average criteria day (E) and average number of trials (F) initiated by the Ts65Dn trisomic mice versus WT littermates. **G-I:** The average percent of mice reaching criteria daily (G), average criteria day (H) and average number of trials (I) initiated by the Ts1Cje trisomic mice versus WT littermates.

**Figure 2:**
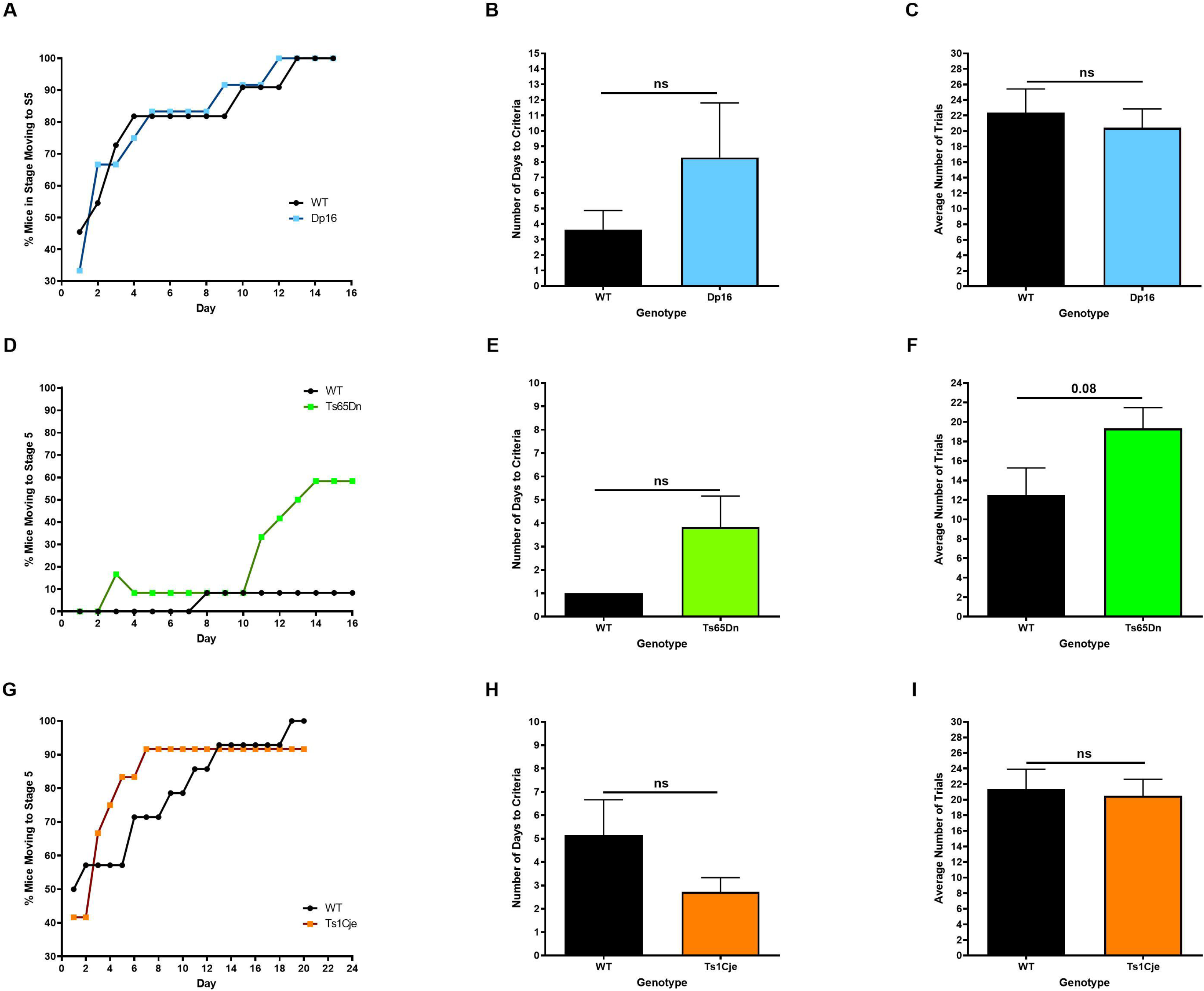
Dp(16)1/Yey, Ts65Dn and Ts1Cje Trisomic Mice Performances in Stage 4 of Pre- Training Compared to WT Littermates. **A-C:** The average percent of mice reaching criteria daily (A), average criteria day (B) and average number of trials (C) initiated by the Dp(16)1/Yey trisomic mice versus WT littermates. **D-F:** The average percent of mice reaching criteria daily (D), average criteria day (E) and average number of trials (F) initiated by the Ts65Dn trisomic mice versus WT littermates. **G-I:** The average percent of mice reaching criteria daily (G), average criteria day (H) and average number of trials (I) initiated by the Ts1Cje trisomic mice versus WT littermates.

All Ts1Cje mice and their WT littermates successfully completed early pre-training at some point throughout the experimental period (Figure 1–2G). There were no differences in the average number of days it took trisomic and WT animals to reach criteria at all Stages of early pre-training (Figure 1–2 H, Table 1). No differences were noted in the average number of trials that were initiated by trisomic and WT mice at all Stages (Figure 1–2 I, Table 1).

##### Late pre-training (Stage 5)

For the Dp(16)1/Yey mice and their WT littermates, there were no differences in the following parameters: average number of days to reach criteria (70% correct answers), percentage of animals reaching criteria each day, number of trials and percentage of correct responses (Figure 3 A-D, Table 1).

**Figure 3:**
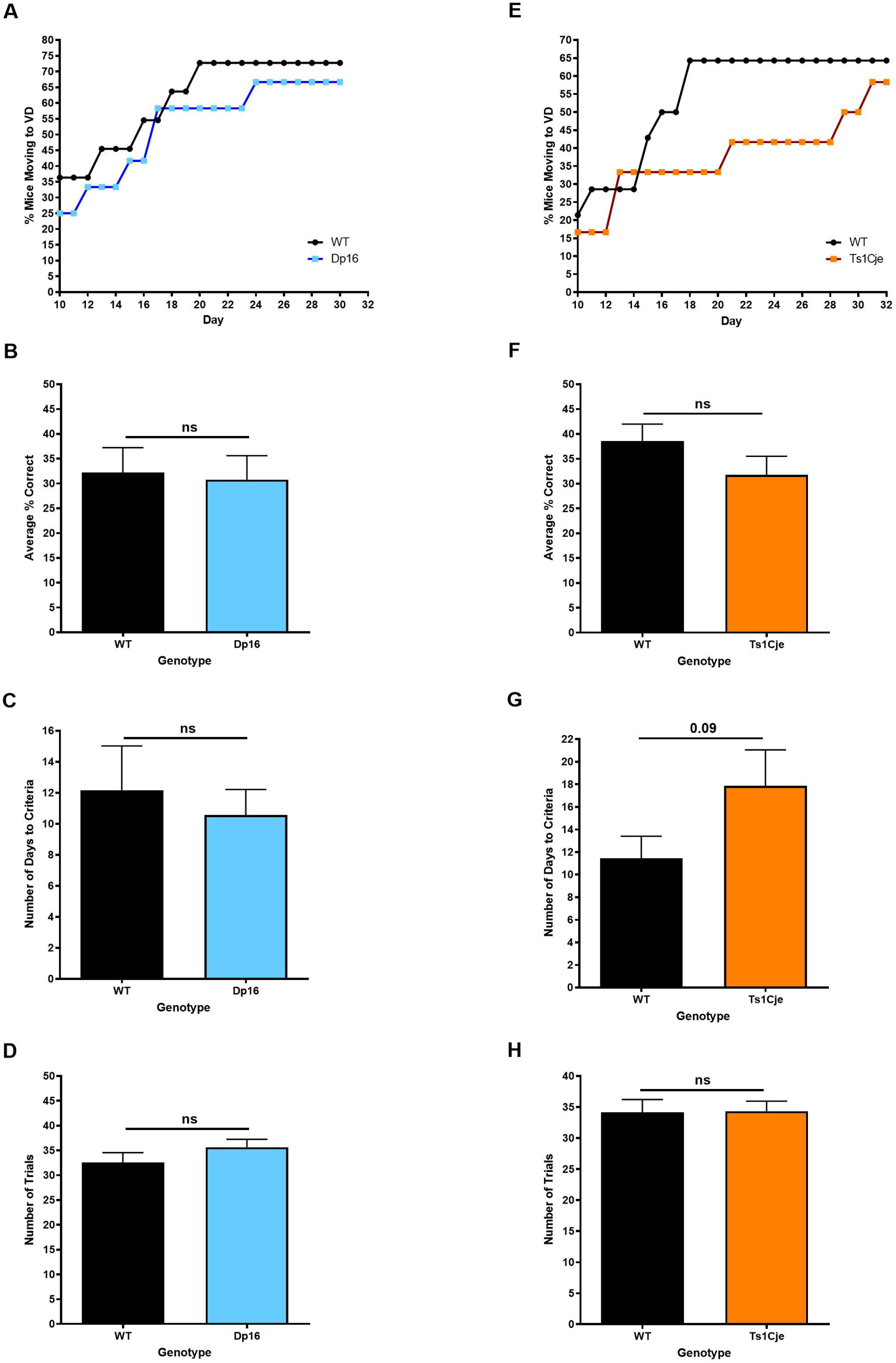
Dp(16)1/Yey and Ts1Cje Trisomic Mice Performances in Stage 5 of Pre-Training Compared to WT Littermates. **A-D:** The average percent of mice reaching criteria daily (A), average percent of correct answers (B), average criteria day (C) and average number of trials (D) initiated by the Dp(16)1/Yey trisomic mice versus WT littermates. **E-H:** The average percent of mice reaching criteria daily (E), average percent of correct answers (F), average criteria day (G) and average number of trials (H) initiated by the Ts1Cje trisomic mice versus WT littermates.

For the Ts65Dn mice and their WT littermates, there were also no differences in the above parameters, however there was only one WT compared to seven Ts65Dn trisomic mice performing at Stage 5 (Data not shown).

For the Ts1Cje mice and their WT littermates, there was a significant difference in the percentage of animals reaching criteria each day, with significantly more WT animals advancing to the Visual Discrimination (VD) Stage each day than Ts1Cje mice (Figure 3E, Table 1). The Ts1Cje mice took more time to reach VD compared to WT littermates, but this difference did not reach statistical significance (*P*=*0.09*) (Figure 3F, Table 1). No differences in the average number of trials or percentage of correct responses were observed (Figure 3EF-H, Table 1).

#### Visual Discrimination

For the Dp(16)1/Yey cohort, there was a significant difference in the average percentage of correct responses between the trisomic and WT littermates (Dp(16)1/Yey =22.70 ± 1.93 %, WT= 32.18 ± 1.49 %, *P*<0.01) (Figure 4A, Table 2). The daily performance of Dp(16)1/Yey trisomic versus WT mice was significantly delayed as indicated by the percent of correct answers every day (Figure 4B, Table 2). However, there was no difference in the average number of trials that were initiated at VD by each genotype (Figure 4C, Table 2).

**Table 2:**
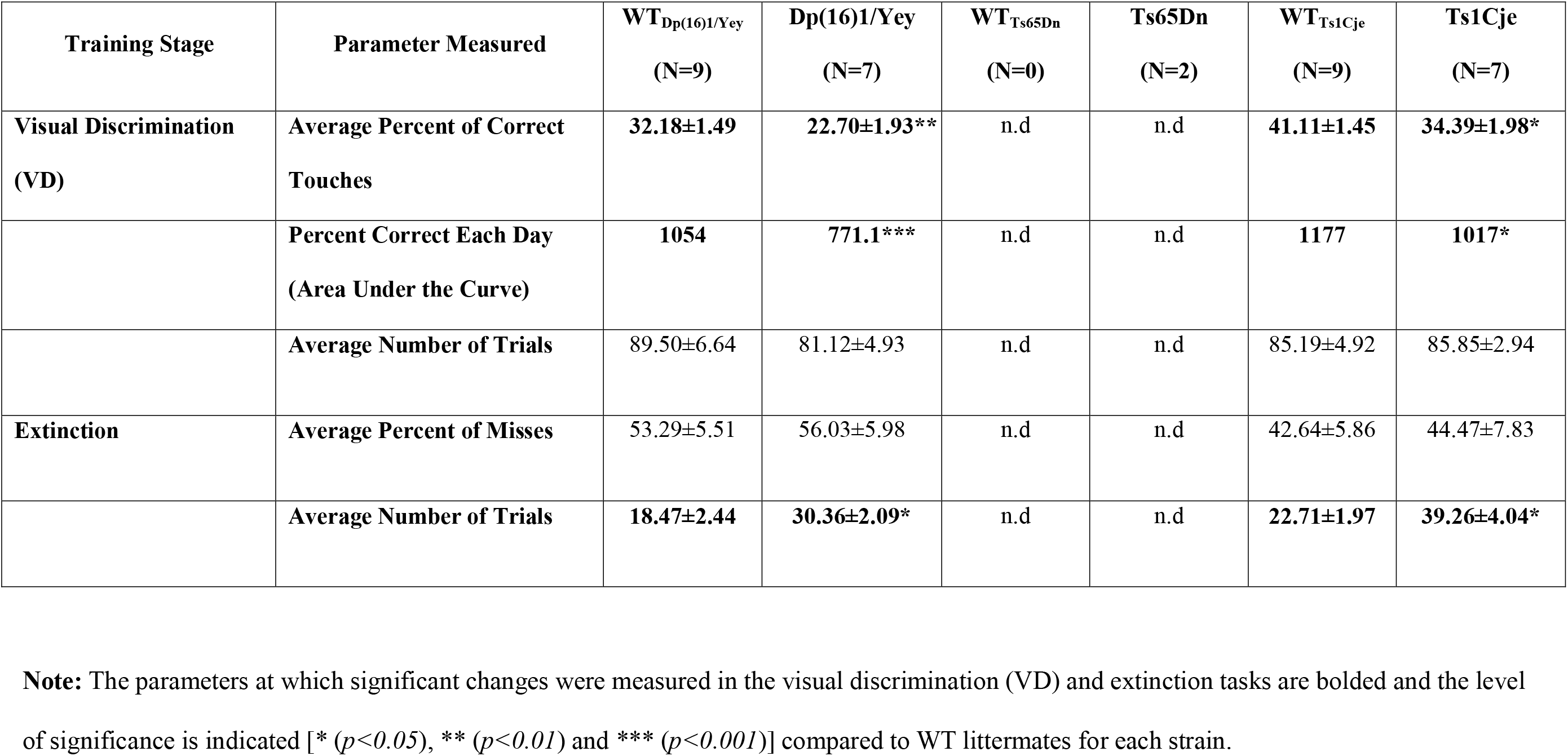
Visual Discrimination (VD) and Extinction Results Summary in the Dp(16)1/Yey, Ts65Dn and Ts1Cje Mouse Models.

| Training Stage | Parameter Measured | WT<sub>Dp(16)1/Yey</sub><br>(N=9) | Dp(16)1/Yey<br>(N=7) | WT<sub>Ts65Dn</sub><br>(N=0) | Ts65Dn<br>(N=2) | WT<sub>Ts1Cje</sub><br>(N=9) | Ts1Cje<br>(N=7) |
| --- | --- | --- | --- | --- | --- | --- | --- |
| **Visual Discrimination**<br>**(VD)** | **Average Percent of Correct**<br>**Touches** | **32.18±1.49** | **22.70±1.93\*\*** | n.d | n.d | **41.11±1.45** | **34.39±1.98\*** |
|  | **Percent Correct Each Day**<br>**(Area Under the Curve)** | **1054** | **771.1\*\*\*** | n.d | n.d | **1177** | **1017\*** |
|  | **Average Number of Trials** | 89.50±6.64 | 81.12±4.93 | n.d | n.d | 85.19±4.92 | 85.85±2.94 |
| **Extinction** | **Average Percent of Misses** | 53.29±5.51 | 56.03±5.98 | n.d | n.d | 42.64±5.86 | 44.47±7.83 |
|  | **Average Number of Trials** | **18.47±2.44** | **30.36±2.09\*** | n.d | n.d | **22.71±1.97** | **39.26±4.04\*** |

**Note:** The parameters at which significant changes were measured in the visual discrimination (VD) and extinction tasks are bolded and the level of significance is indicated [\* ( $p<0.05$ ), \*\* ( $p<0.01$ ) and \*\*\* ( $p<0.001$ )] compared to WT littermates for each strain.

**Figure 4:**
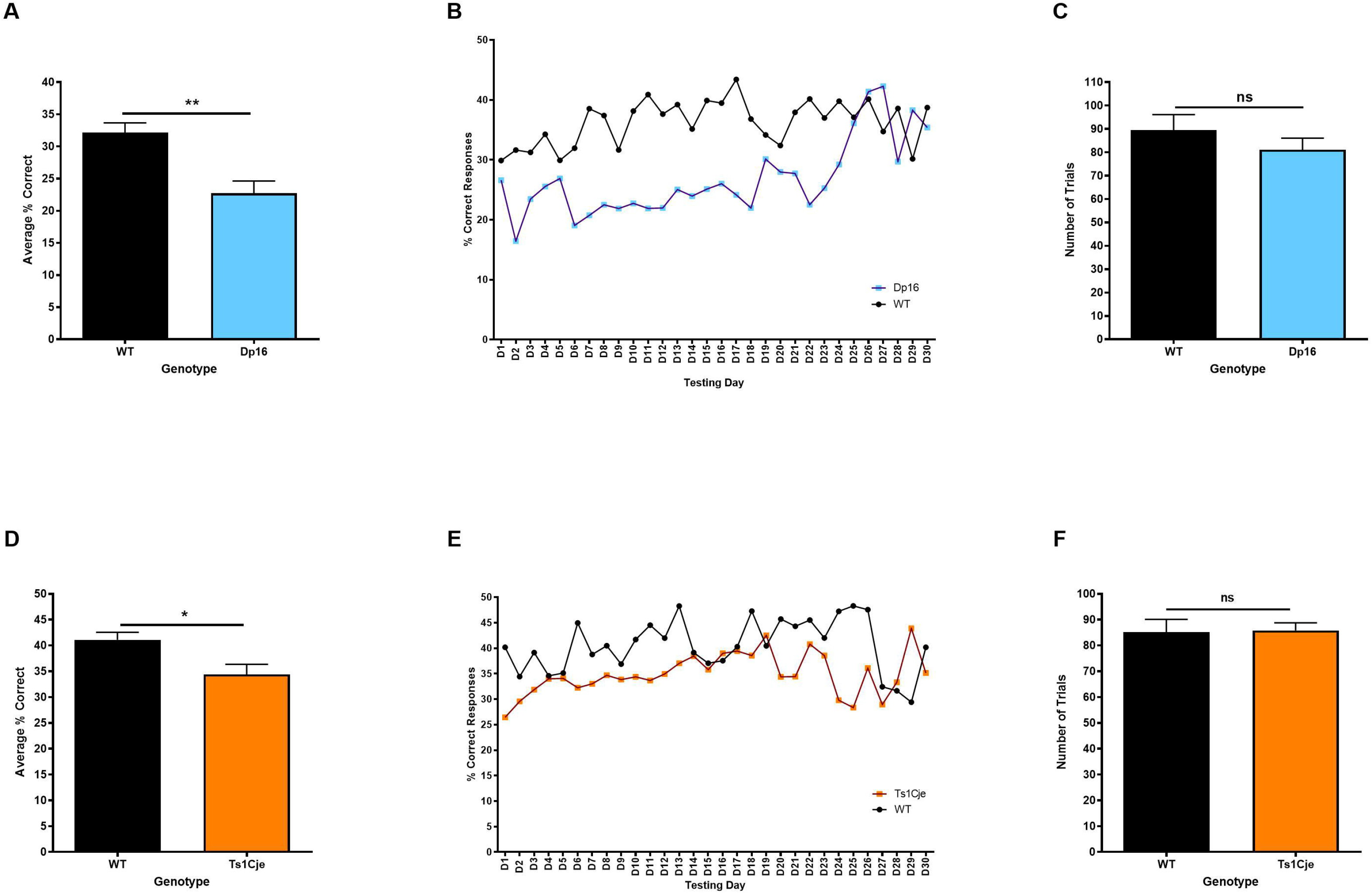
Dp(16)1/Yey and Ts1Cje Trisomic Mice Performances in the Visual Discrimination Task Compared to WT Littermates. **A-C:** The average percent of correct responses (A), the percent of correct answers each day (B) and the average number of trials (D) initiated by the Dp(16)1/Yey trisomic mice versus WT littermates. **D-F:** The average percent of correct responses (D), the percent of correct answers each day (E) and the average number of trials (F) initiated by the Ts1Cje trisomic mice versus WT littermates.

For the Ts65Dn strain, among the seven Ts65Dn mice and one WT littermate that were tested in Stage 5 of pre-training, only two Ts65Dn mice were able to reach VD. Thus, no VD data were analyzed for the Ts65Dn cohort.

The Ts1Cje cohort, there was a significant difference in the average percentage of correct responses between the Ts1Cje and WT animals (Ts1Cje=34.39 ± 1.98 %, WT= 41.11 ± 1.45 %, *P*<0.05) (Figure, Table 2). Although similar findings were observed in the Ts1Cje and Dp(16)1/Yey trisomic mice, the latter exhibited more severe learning deficits in the VD task (Figure 4 B, E, Table 2). There were no differences in the average number of trials initiated by all genotypes.

#### Extinction

For the Dp(16)1/Yey cohort, the average percent of misses were not significantly different between the trisomic and WT mice (Figure 5A, Table 2). However, Dp(16)1/Yey mice initiated a significantly higher number trials (30.36 ± 2.08) compared to their WT littermates (18.47 ± 2.44) (*P*<0.01) (Figure 5 B-C, Table 2).

**Figure 5:**
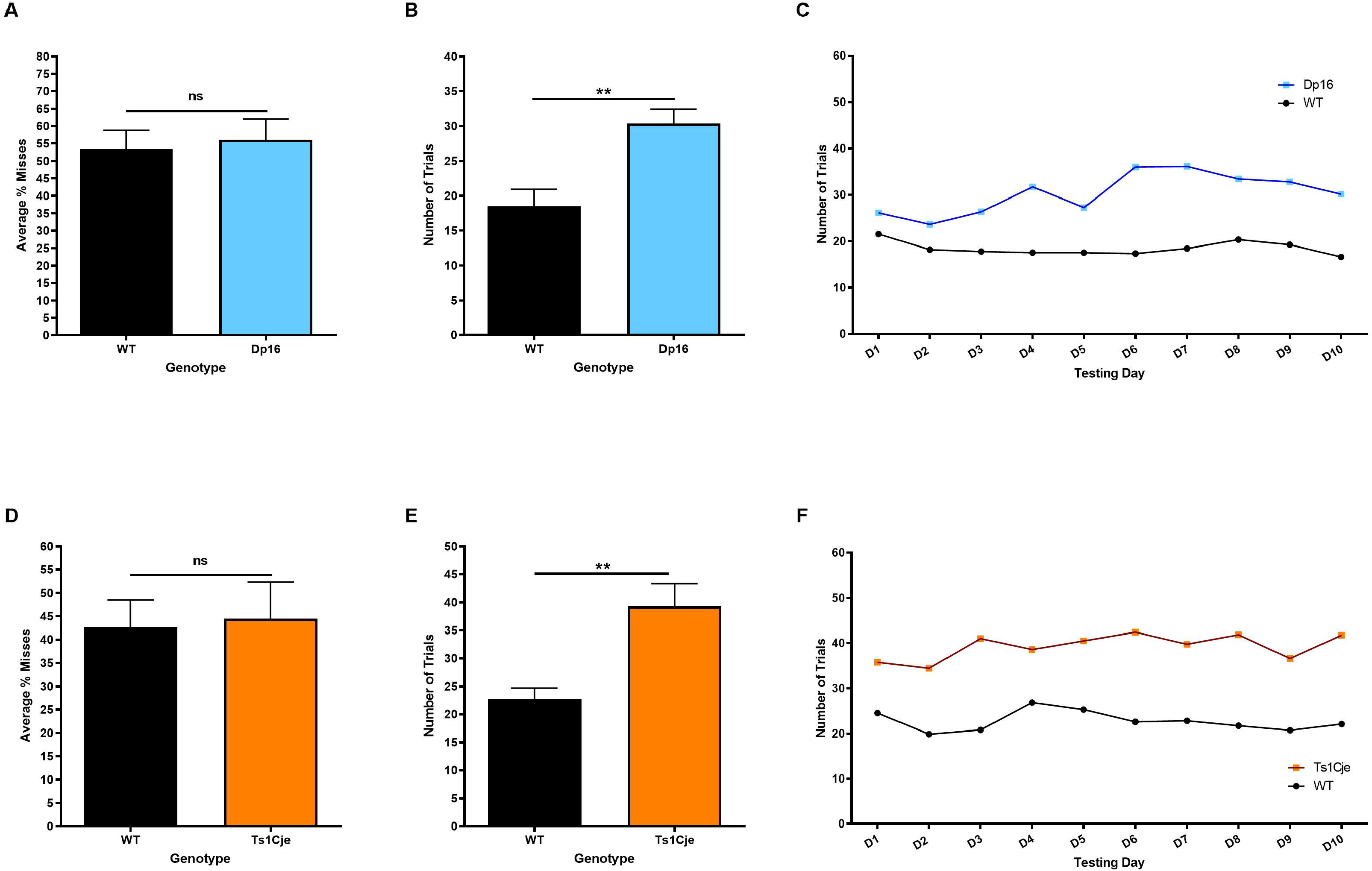
Dp(16)1/Yey and Ts1Cje Trisomic Mice Performances in the Extinction Task Compared to WT Littermates. **A-C:** The average percent of misses (A), the average number of trials (B) and the number of trials performed each day (C) initiated by the Dp(16)1/Yey trisomic mice versus WT littermates. **D-F:** The average percent of misses (D), the average number of trials (E) and the number of trials performed each day (F) initiated by the Ts1Cje trisomic mice versus WT littermates.

Similar to what had been observed in the Dp(16)1/Yey cohort, there was no difference in the average percent of misses between Ts1Cje and WT littermate mice (Figure 5D, Table 2), but there was also a significant difference in the number of trials initiated by Ts1Cje mice (39.26 ± 4.04) compared to their WT littermates (22.71 ± 1.97, p<0.001) (Figure 5E-F, Table 2).

No Ts65Dn mice were tested in the extinction phase because most mice did not reach the VD Stage.

### II The effects of mild food restriction on training performance in C57BL/6J mice

In response to the prolonged time that it took trisomic and WT littermates to reach criterion at most Stages of our operant conditioning experiment, we tested a cohort of five-week-old, naïve C57BL/6J male mice (N=10) under the same protocols, five of which underwent food restriction. The goal was to determine whether mild food restriction (15 % reduction of body weight) would decrease the amount of time it would take the mice to advance to subsequent Stages in our protocol, as well as increase the number of correct touches and number of trials initiated per session. The results showed that all five food-restricted mice reached criteria at all Stages and in significantly less days than their non-restricted counterparts, of which only two advanced beyond Stage 2 (Figure 6 A, D).

**Figure 6:**
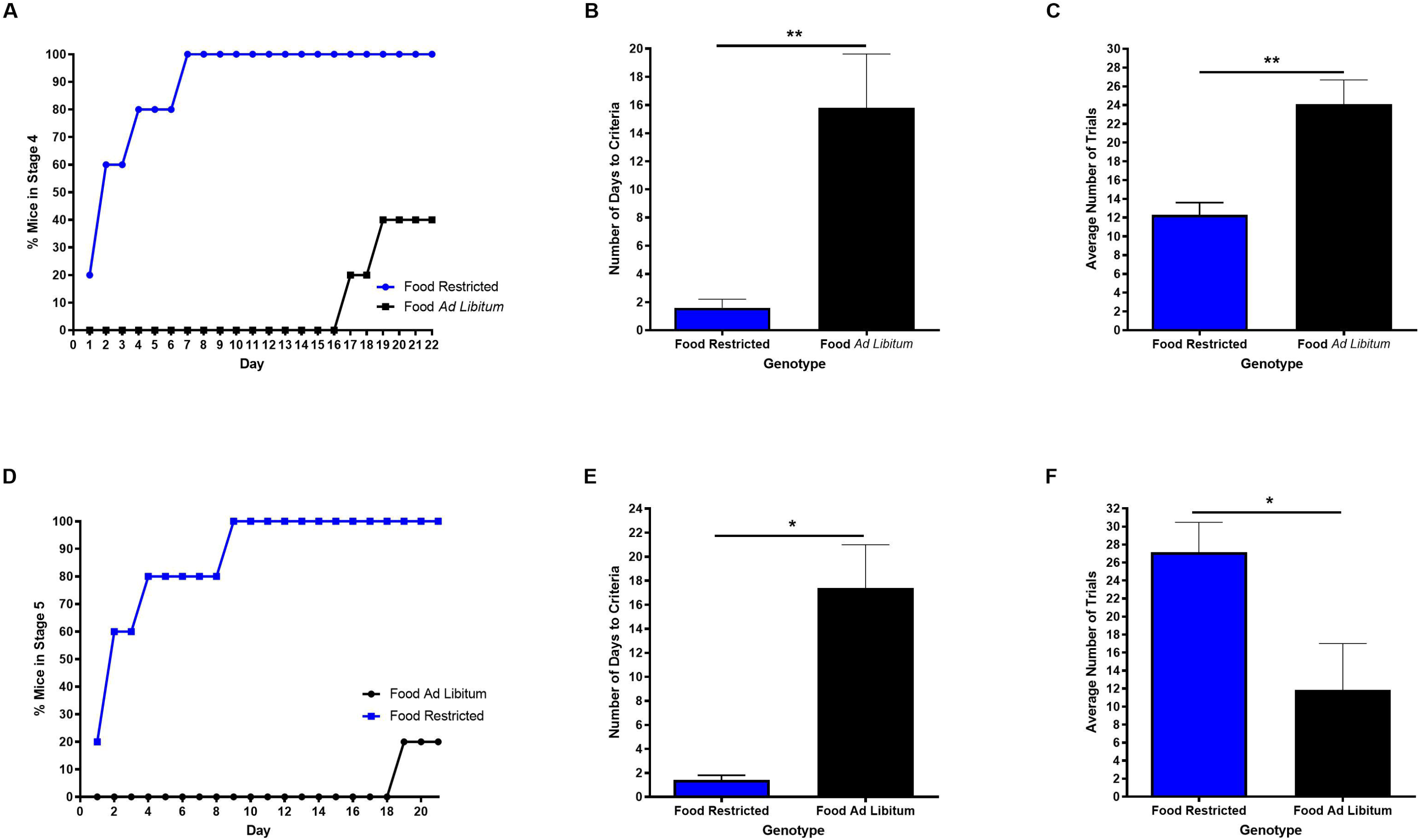
Effects of Mild Food Restriction on Pre-Training Performance in C57BL/6J mice. **A-C:** The average percent of mice reaching criteria daily (A), average criteria day (B) and average number of trials (C) initiated in pre-training Stage 3 by food restricted C57BL/6J mice compared to C57BL/6J that had *ad libitum* access to food and water. **D-F:** The average percent of mice reaching criteria daily (A), average criteria day (B) and average number of trials (C) initiated in pre-training Stage 4 by the food restricted C57BL/6J mice compared to C57BL/6J that had *ad libitum* access to food and water.

#### Pre-training

##### Early pre-training (Stages 2-4)

The average number of days to reach criteria, as well as the percentage of animals advancing to the next Stage each day were significantly different between restricted and non-restricted mice at all Stages of early pre-training (Figure 6 A, D, Table 3). As a result, there was a significant reduction in the average number of days to reach criteria in the food restricted C57BL/6J compared to mice that had access to food and water *ad libitum* (Figure 6 B, E, Table 3). There were also significant differences in the average number of trials initiated at Stages 3 and 4 between both cohorts with food restricted mice performing significantly more trials than their non-restricted counterparts (Figure 6 C, F, Table 3).

**Table 3:**
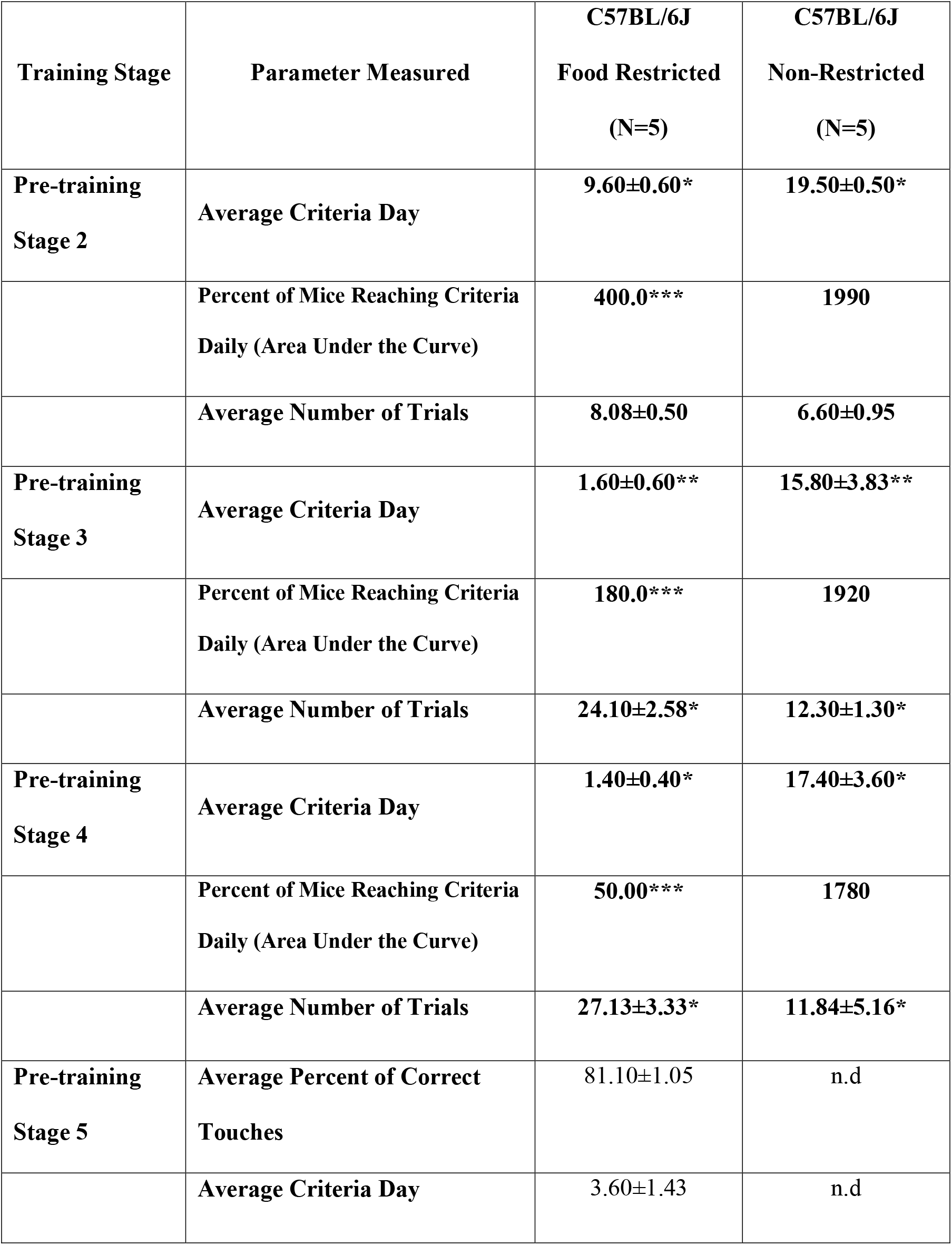
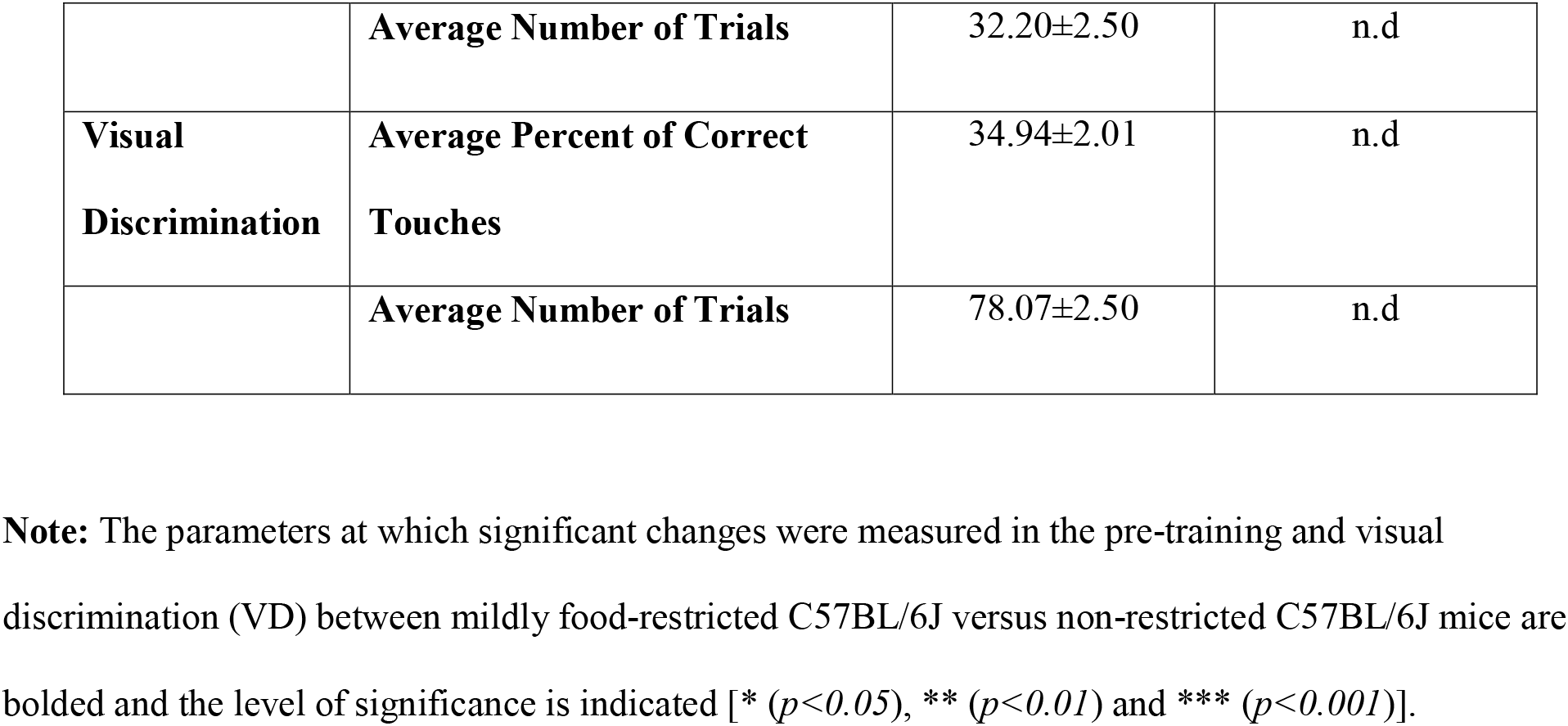
Pre-Training and Visual Discrimination Results Summary in food restricted versus non-restricted C56BL/6J mice.

##### Late pre-training (Stage 5)

The average number of days to reach criteria was 3.60 ± 1.43 days for the restricted mice (Figure 6 E, Table 3). Only one non-restricted C57BL/6J mouse advanced from Stage 4 to Stage 5, thus no statistical analysis was performed at Stage 5 for the average number of days to reach criteria, the percent of correct answers and the average number of trials. These data are presented in Table 3 for the food restricted C57BL/6J mice.

##### Visual Discrimination (VD)

All five food restricted C57BL/6J mice advanced to VD and no non-restricted mice reached this Stage, thus the average number of trials and the percent of correct answers were only collected for the food restricted C57BL/6J mice (Table 3). The average number of trials performed by the food restricted C57BL/6J mice was 78.07 ± 2.50 trials/session with a percent of correct answers averaging 34.94 ± 2.01 % (Table 3).

## Discussion

Down syndrome associated intellectual disability has been the target of many pre-clinical and human clinical trials to improve cognition and independent life skills ^3–5^. These major efforts were largely unsuccessful due to several reasons, including the choice of the best mouse models, the strategies used for drug development, the timing of intervention and the lack of human translational cognitive outcome measures in preclinical studies ^7^. Although conventional rodent behavioral tests offer a critical and quick way to evaluate learning and memory deficits in mouse models of central nervous system (CNS) disorders and to evaluate the efficacy of treatments, these behavioral paradigms do not translate to humans. Data obtained from them might not directly reflect the potential effects of these treatments in human clinical trials ^19,20^. Recently, rodent touchscreen based behavioral platforms have emerged as an alternative for conventional rodent behavioral tests. These touchscreen platforms use technologies and cognitive tests that were directly adapted from human clinical studied to narrow the translational gap between pre- clinical and clinical trials ^21^.

In this study, we used touchscreen based behavioral testing to investigate hippocampal-dependent learning and memory (Visual Discrimination task) and frontal lobe inhibitory control (Extinction task) in three cytogenetically different mouse models of DS, including Dp(16)1/Yey, Ts65Dn and Ts1Cje mouse models. No aversive stimuli (food or water restriction) were used in our study in order to be able to mix potential treatments with food in future studies and to ensure correct dosage and intake of the drug molecules by treated animals. Pre-peer review data from our laboratory has demonstrated that one of these promising drug candidates, namely apigenin, showed improvement of neonatal olfactory spatial memory and adult hippocampal-dependent memory in the Ts1Cje mouse model of Down syndrome ^22^.

### I Performances in the Dp(16)1/Yey, T65Dn and Ts1Cje mouse models

#### Deficits During the Pre-training Stage

During early pre-training phases (Stages 2 to 4), Dp(16)1/Yey and Ts1Cje trisomic mice did not show any learning/memory deficits and progressed at the same rate to late pre-training (Stage 5) compared to their WT littermates. During late pre-training, Dp(16)1/Yey trisomic and WT littermate mice exhibited similar performances and reached VD at the same rate while Ts1Cje trisomic mice showed a significant delay in advancing to VD.

Somewhat unexpectedly, Ts65Dn trisomic mice outperformed their WT littermates during early pre-training. A significantly higher percentage of Ts65Dn trisomic mice advanced through the pre-training Stages compared to WT littermates. Notably, this faster advancement of the Ts65Dn trisomic mice during the pre-training Stages is likely due to the significant increase in the number of trials they initiated compared to WT littermates rather than faster learning. This observation is supported by our previous study in which we demonstrated that Ts65Dn trisomic mice exhibited hyperactivity versus WT littermates in the open field test, which might explain their increased number of trials in the present study ^18^. Additionally, it is well documented in multiple studies that Ts65Dn trisomic mice exhibit significant hippocampal-dependent learning and memory deficits compared to WT littermates ^18, 23–25^.

To our knowledge, only one study investigated hippocampal-dependent memory in food-restricted Ts65Dn mice using a pairwise Visual Discrimination task ^26^. Similar to our findings here, Ts65Dn mice advanced through pre-training faster than WT littermates, however, this study demonstrated a mild delay in the number of days but a significant increase in the number of trials to reach criteria compared to WT littermates.

In our study, very few Ts65Dn trisomic mice and their WT littermates advanced successfully to pre-training Stage 5 and VD compared to the Dp(16)1/Yey and Ts1Cje strains. Differences in the genetic background between the Ts65Dn strain and the other two mouse models (i.e. Dp(16)1/Yey and Ts1Cje) might be responsible for the differences in touchscreen performances observed in non-food restricted conditions. Indeed, the Ts65Dn mice were maintained on an F1 hybrid background (C57BL/6J x C3H) while Dp(16)1/Yey and Ts1Cje mice were maintained on a C57BL/6J background.

Unfortunately, there are no literature reports describing back-crossing of the Ts65Dn mouse model onto a C57BL/6J and their performances in different behavioral paradigms.

A closer examination of the data by Leach and Crawley (2018) ^26^ shows clearly that Ts65Dn and their WT littermates maintained on an F1 hybrid genetic background required significantly more time (over 60 days for Ts65Dn and 40 days for WT littermates) and performed significantly more trials (close to 4000 trials for Ts65Dn and over 2000 trials for WT littermates) to reach criteria compared to the *Ube3a* knock-out mice (less than 30 days and 1500 trials) and their WT littermates (less than 15 days and less than 1000 trials) that were maintained on a C57BL/6J genetic background. These data suggest that the genetic background of the Ts65Dn mice significantly impacts their performances in touchscreen-based learning/memory tasks.

Several studies highlighted strain background differences in learning/memory, circadian rhythm and aging in different inbred strains ^27–31^. Interestingly, Brooks et al (2005) ^32^ analyzed the learning/memory profiles of several inbred mouse strains using different cognitive tasks, including the non-touchscreen based Visual Discrimination (VD) test and Morris water maze. They demonstrated that C3H mice were unable to acquire the VD task and showed lower performances in the Morris water maze compared to the other strains.

#### Deficits During the Visual Discrimination Task

Both the Dp(16)1/Yey and Ts1Cje trisomic mice showed significant deficits in their VD performances versus WT littermates without differences in the average number of trials performed by each genotype. It is worth noting that the Dp(16)1/Yey exhibited more severe learning deficits than the Ts1Cje strain. These findings confirm our previously published data using conventional rodent behavioral tests that showed very severe hippocampal-dependent contextual (fear conditioning test) and spatial memory (Morris water maze test) in the Dp(16)1/Yey trisomic mice. Using those same tests, we also showed that the Ts1Cje mice have milder hippocampal-dependent memory deficits compared to the Dp(16)1/Yey and Ts65Dn mice^18^.

Several cognitive batteries have been used to extensively study language, executive function, learning and memory as well as attention and psychomotor speed in humans with DS. Among these testing batteries, several CANTAB tests have been used to investigate the above-mentioned domains, including Visual Pattern Recognition (or Visual Discrimination), Paired Associates Learning (PAL), Reaction Time (RTI), Spatial Recognition (SR) and spatial span (SSP) ^14,17, 33–35^.

Individuals with DS exhibit significant hippocampal-dependent learning/memory deficits as demonstrated by lower response accuracy in the CANTAB Visual Pattern Recognition test as well as a reduction in the number of correct responses and total number of Stages reached in the CANTAB PAL task compared to mental age matched typically developing individuals ^15–16,36^. However, individuals with DS and their mental age matched controls performed similarly in the prefrontal cortex-dependent cognitive tasks like the CANTAB Stockings of Cambridge ^36^.

#### Deficits During the Extinction Task

During the Extinction Stage, both Dp(16)1/Yey and Ts1Cje trisomic mice had similar average and daily percent of misses compared to their WT littermates. However, they performed a significantly higher number of trials on a daily basis and overall. The increased number of trials suggests that both Dp(16)1/Yey and Ts1Cje mice exhibit a perseverative behavior, repetitive behavior and/or attention-deficit-hyperactivity disorder. Similar findings were reported for the *mGluR5* (metabotropic glutamate receptor 5) knock-out mouse model ^37,38^.

Children with attention-deficit-hyperactivity disorder (ADHD) exhibited normal hippocampal-dependent learning (Visual Discrimination) but significant delays in the learning flexibility (Rule Reversal) and extinction learning compared to typically developing children ^39^.

Although extinction learning is understudied in DS, several studies demonstrated that humans with DS have high prevalence of ADHD (34%) and autism spectrum disorder or ASD (16-42%) ^40–42^.

### II The effects of mild food restriction on training performance in C57BL/6J mice

To increase animal motivation and reduce the time needed to complete the touchscreen behavior tasks, we compared performances of C57BL/6J mice under food restriction with a second group of C57BL/6J mice having *ad libitum* access to food and water. At all pre-training Stages, food restricted C57BL/6J animals outperformed non-restricted C57BL/6J with a significant reduction in the number of days needed to reach criteria along with increased number of trials performed and the percent of correct answers, thus significantly reducing the pre-training duration to less than two weeks.

Multiple studies suggested that a food restriction resulting in up to 15% body weight reduction does not significantly affected the physiological parameters in a rodent ^43,44^. Comparison of the motivational effects of food and water restriction in a fixed-head Visual Discrimination task demonstrated that mice trained under food-restriction executed significantly more trials than water-restricted mice during the pre-training and testing Stages, however, water-restricted mice reached criteria faster in the VD task ^45^. In another study, Forestell et al (2001) ^46^ investigated the performances of food restricted CD1 mice in an odor discrimination task and showed that these mice had higher motivation and faster learning compared to mice that had *ad libitum* access to food. In another study, Tucci et al (2006) compared the effects of food and water restriction in C57BL/6J on motivation and learning in a nose poking operant learning task. Both food and water restricted mice showed higher peak of responses compared non-restricted mice ^47^.

Although aversive stimuli such as food and water restriction strongly increase animal motivation to perform operant and touchscreen-based tasks, future studies are needed to investigate their impact on drug intake, dosage and pharmacokinetics in preclinical therapeutic trials of CNS disorders.

## Conclusions

In the present study, we were able to successfully translate the touchscreen-based CANTAB Visual Discrimination and Extinction tasks to evaluate hippocampal-dependent learning and frontal lobe-dependent inhibitory control in the three most commonly used mouse models of DS, including Dp(16)1/Yey, Ts65Dn and Ts1Cje models. We demonstrated that Dp(16)1/Yey trisomic mice exhibited the most severe delay in hippocampal-dependent learning whereas Ts1Cje mice showed a milder delay. Ts65Dn mice outperformed their WT littermates during pre-training but both genotypes were unable to reach the VD Stage. In the Extinction task, both Dp(16)1/Yey and Ts1Cje performed significantly more trials than their WT littermates suggesting that hyperactive and compulsive behaviors are present in these two models. Finally, in a separate set of experiments, we demonstrated that mild food restriction (85% of body weight) significantly increased animal motivation and reduced the time required to complete the touchscreen-based testing.

## Future Studies

Neuroimaging studies have reported region-specific hypoplasia of the frontal lobe, hippocampus and cerebellum in humans with DS. These neuroanatomical defects are associated with major deficits in the cognitive and motor domains linked to those brain regions.

Future studies will examine the effects of mild food restriction on these cognitive and motor domains in the Dp(16)1/Yey, Ts65Dn and Ts1Cje mouse models using several CANTAB based tasks targeting hippocampal-dependent learning/memory (Visual Discrimination and Paired Associates Learning), hippocampal-dependent spatial memory (Trial Unique Delayed Non-match to Location) cortical-dependent learning flexibility (Rule Reversal), inhibitory control (Extinction) and cerebellum-dependent motor learning (5 Choice Serial Reaction Time task).

The goal of these studies is to identify the best mouse model that replicates the human DS cognitive and motor phenotypes and that can be used for preclinical treatment studies.

We will then use the most affected cognitive domains as endpoints to evaluate the effects of prenatal and/or postnatal therapeutic interventions in combination with the neuroanatomical, molecular, cellular and behavioral endpoints identified in our previous studies ^18,48^.

## Acknowledgments

This work was funded by the National Institutes of Health (NICHD R01HD058880) and the National Human Genome Research Institute’s Intramural Research Program (NHGRI Z1A HG200399-04). The funders had no role in study design, data collection and analysis, decision to publish, or preparation of the manuscript.

## Competing Interests

No competing interests declared.

## Notes

### Competing Interest Statement

The authors have declared no competing interest.

## References

1. Delabar, J.M., Aflalo-Rattenbac, R. & Creau, N. (2006) Developmental defects in trisomy 21 and mouse models. ScientificWorldJournal, 6, 1945–1964.

2. Delabar, J.M., Allinquant, B., Bianchi, D., Blumenthal, T., Dekker, A., Edgin, J., O’Bryan, J., Dierssen, M., Potier, M.C., Wiseman, F., Guedj, F., Creau, N., Reeves, R., Gardiner, K. & Busciglio, J. (2016) Changing Paradigms in Down Syndrome: The First International Conference of the Trisomy 21 Research Society. Mol Syndromol, 7, 251–261.

3. Guedj, F., Bianchi, D.W. & Delabar, J.M. (2014) Prenatal treatment of Down syndrome: a reality? Curr Opin Obstet Gynecol, 26, 92–103.

4. Stagni, F., Giacomini, A., Guidi, S., Ciani, E. & Bartesaghi, R. (2015) Timing of therapies for Down syndrome: the sooner, the better. Front Behav Neurosci, 9, 265.

5. Costa, A.C. & Scott-McKean, J.J. (2013) Prospects for improving brain function in individuals with Down syndrome. CNS Drugs, 27, 679–702.

6. Hart, S.J., Visootsak, J., Tamburri, P., Phuong, P., Baumer, N., Hernandez, M.C., Skotko, B.G., Ochoa-Lubinoff, C., Liogier D’Ardhuy, X., Kishnani, P.S. & Spiridigliozzi, G.A. (2017) Pharmacological interventions to improve cognition and adaptive functioning in Down syndrome: Strides to date. Am J Med Genet A, 173, 3029–3041.

7. Lee, S.E., Duran-Martinez, M., Khantsis, S., Bianchi, D.W. & Guedj, F. (2020) Challenges and Opportunities for Translation of Therapies to Improve Cognition in Down Syndrome. Trends Mol Med, 26, 150–169.

8. De la Torre, R., De Sola, S., Pons, M., Duchon, A., de Lagran, M.M., Farre, M., Fito, M., Benejam, B., Langohr, K., Rodriguez, J., Pujadas, M., Bizot, J.C., Cuenca, A., Janel, N., Catuara, S., Covas, M.I., Blehaut, H., Herault, Y., Delabar, J.M. & Dierssen, M. (2014) Epigallocatechin-3-gallate, a DYRK1A inhibitor, rescues cognitive deficits in Down syndrome mouse models and in humans. Mol Nutr Food Res, 58, 278–288.

9. de la Torre, R., de Sola, S., Hernandez, G., Farre, M., Pujol, J., Rodriguez, J., Espadaler, J.M., Langohr, K., Cuenca-Royo, A., Principe, A., Xicota, L., Janel, N., Catuara-Solarz, S., Sanchez-Benavides, G., Blehaut, H., Duenas-Espin, I., Del Hoyo, L., Benejam, B., Blanco-Hinojo, L., Videla, S., Fito, M., Delabar, J.M., Dierssen, M. & group, T.s. (2016) Safety and efficacy of cognitive training plus epigallocatechin-3-gallate in young adults with Down’s syndrome (TESDAD): a double-blind, randomised, placebo-controlled, phase 2 trial. Lancet Neurol, 15, 801–810.

10. Sahakian, B.J. & Owen, A.M. (1992) Computerized assessment in neuropsychiatry using CANTAB: discussion paper. J R Soc Med, 85, 399–402.

11. Weintraub, S., Bauer, P.J., Zelazo, P.D., Wallner-Allen, K., Dikmen, S.S., Heaton, R.K., Tulsky, D.S., Slotkin, J., Blitz, D.L., Carlozzi, N.E., Havlik, R.J., Beaumont, J.L., Mungas, D., Manly, J.J., Borosh, B.G., Nowinski, C.J. & Gershon, R.C. (2013) I. NIH Toolbox Cognition Battery (CB): introduction and pediatric data. Monogr Soc Res Child Dev, 78, 1–15.

12. Akshoomoff, N., Newman, E., Thompson, W.K., McCabe, C., Bloss, C.S., Chang, L., Amaral, D.G., Casey, B.J., Ernst, T.M., Frazier, J.A., Gruen, J.R., Kaufmann, W.E., Kenet, T., Kennedy, D.N., Libiger, O., Mostofsky, S., Murray, S.S., Sowell, E.R., Schork, N., Dale, A.M. & Jernigan, T.L. (2014) The NIH Toolbox Cognition Battery: results from a large normative developmental sample (PING). Neuropsychology, 28, 1–10.

13. Hessl, D., Sansone, S.M., Berry-Kravis, E., Riley, K., Widaman, K.F., Abbeduto, L., Schneider, A., Coleman, J., Oaklander, D., Rhodes, K.C. & Gershon, R.C. (2016) The NIH Toolbox Cognitive Battery for intellectual disabilities: three preliminary studies and future directions. J Neurodev Disord, 8, 35.

14. Esbensen, A.J., Hooper, S.R., Fidler, D., Hartley, S.L., Edgin, J., d’Ardhuy, X.L., Capone, G., Conners, F.A., Mervis, C.B., Abbeduto, L., Rafii, M.S., Krinsky-McHale, S.J., Urv, T. & Outcome Measures Working, G. (2017) Outcome Measures for Clinical Trials in Down Syndrome. Am J Intellect Dev Disabil, 122, 247–281.

15. Visu-Petra, L., Benga, O., Tincas, I. & Miclea, M. (2007) Visual-spatial processing in children and adolescents with Down’s syndrome: a computerized assessment of memory skills. J Intellect Disabil Res, 51, 942–952.

16. Edgin, J.O., Mason, G.M., Allman, M.J., Capone, G.T., Deleon, I., Maslen, C., Reeves, R.H., Sherman, S.L. & Nadel, L. (2010) Development and validation of the Arizona Cognitive Test Battery for Down syndrome. J Neurodev Disord, 2, 149–164.

17. Edgin, J.O., Anand, P., Rosser, T., Pierpont, E.I., Figueroa, C., Hamilton, D., Huddleston, L., Mason, G., Spano, G., Toole, L., Nguyen-Driver, M., Capone, G., Abbeduto, L., Maslen, C., Reeves, R.H. & Sherman, S. (2017) The Arizona Cognitive Test Battery for Down Syndrome: Test-Retest Reliability and Practice Effects. Am J Intellect Dev Disabil, 122, 215–234.

18. Aziz, N.M., Guedj, F., Pennings, J.L.A., Olmos-Serrano, J.L., Siegel, A., Haydar, T.F. & Bianchi, D.W. (2018) Lifespan analysis of brain development, gene expression and behavioral phenotypes in the Ts1Cje, Ts65Dn and Dp(16)1/Yey mouse models of Down syndrome. Dis Model Mech, 11.

19. Hvoslef-Eide, M., Nilsson, S.R., Saksida, L.M. & Bussey, T.J. (2016) Cognitive Translation Using the Rodent Touchscreen Testing Approach. Curr Top Behav Neurosci, 28, 423–447.

20. Kangas, B.D. & Bergman, J. (2017) Touchscreen technology in the study of cognition-related behavior. Behav Pharmacol, 28, 623–629.

21. Horner, A.E., Heath, C.J., Hvoslef-Eide, M., Kent, B.A., Kim, C.H., Nilsson, S.R., Alsio, J., Oomen, C.A., Holmes, A., Saksida, L.M. & Bussey, T.J. (2013) The touchscreen operant platform for testing learning and memory in rats and mice. Nat Protoc, 8, 1961–1984.

22. Guedj, F., Pennings, J.LA., Siegel, A.E., Alsebaa, F., Massingham, L.J., Tantravahi, U., Bianchi, D.W (2018). Apigenin as a Candidate Prenatal Treatment for Trisomy 21: Effects in Human Amniocytes and the Ts1Cje Mouse Model. bioRxiv 495283

23. Costa, A.C., Stasko, M.R., Schmidt, C. & Davisson, M.T. (2010) Behavioral validation of the Ts65Dn mouse model for Down syndrome of a genetic background free of the retinal degeneration mutation Pde6b(rd1). Behav Brain Res, 206, 52–62.

24. Martinez-Cue, C., Rueda, N., Garcia, E., Davisson, M.T., Schmidt, C. & Florez, J. (2005) Behavioral, cognitive and biochemical responses to different environmental conditions in male Ts65Dn mice, a model of Down syndrome. Behav Brain Res, 163, 174–185.

25. Escorihuela, R.M., Fernandez-Teruel, A., Vallina, I.F., Baamonde, C., Lumbreras, M.A., Dierssen, M., Tobena, A. & Florez, J. (1995) A behavioral assessment of Ts65Dn mice: a putative Down syndrome model. Neurosci Lett, 199, 143–146.

26. Leach, P.T. & Crawley, J.N. (2018) Touchscreen learning deficits in Ube3a, Ts65Dn and Mecp2 mouse models of neurodevelopmental disorders with intellectual disabilities. Genes Brain Behav, 17, e12452.

27. Lad, H.V., Liu, L., Paya-Cano, J.L., Parsons, M.J., Kember, R., Fernandes, C. & Schalkwyk, L.C. (2010) Behavioural battery testing: evaluation and behavioural outcomes in 8 inbred mouse strains. Physiol Behav, 99, 301–316.

28. Moy, S.S., Nadler, J.J., Young, N.B., Perez, A., Holloway, L.P., Barbaro, R.P., Barbaro, J.R., Wilson, L.M., Threadgill, D.W., Lauder, J.M., Magnuson, T.R. & Crawley, J.N. (2007) Mouse behavioral tasks relevant to autism: phenotypes of 10 inbred strains. Behav Brain Res, 176, 4–20.

29. Banks, G., Heise, I., Starbuck, B., Osborne, T., Wisby, L., Potter, P., Jackson, I.J., Foster, R.G., Peirson, S.N. & Nolan, P.M. (2015) Genetic background influences age-related decline in visual and nonvisual retinal responses, circadian rhythms, and sleep. Neurobiol Aging, 36, 380–393.

30. Wilking, J.A., Hesterberg, K.G., Nguyen, V.H., Cyboron, A.P., Hua, A.Y. & Stitzel, J.A. (2012) Comparison of nicotine oral consumption and baseline anxiety measures in adolescent and adult C57BL/6J and C3H/Ibg mice. Behav Brain Res, 233, 280–287.

31. Logan, R.W., McCulley, W.D., 3rd, Seggio, J.A. & Rosenwasser, A.M. (2012) Effects of withdrawal from chronic intermittent ethanol vapor on the level and circadian periodicity of running-wheel activity in C57BL/6J and C3H/HeJ mice. Alcohol Clin Exp Res, 36, 467–476.

32. Brooks, S.P., Pask, T., Jones, L. & Dunnett, S.B. (2005) Behavioural profiles of inbred mouse strains used as transgenic backgrounds. II: cognitive tests. Genes Brain Behav, 4, 307–317.

33. Vicari, S., Caselli, M.C., Gagliardi, C., Tonucci, F. & Volterra, V. (2002) Language acquisition in special populations: a comparison between Down and Williams syndromes. Neuropsychologia, 40, 2461–2470.

34. Wishart, J.G. (1995) Cognitive abilities in children with Down syndrome: developmental instability and motivational deficits. Prog Clin Biol Res, 393, 57–91.

35. Liogier d’Ardhuy, X., Edgin, J.O., Bouis, C., de Sola, S., Goeldner, C., Kishnani, P., Noldeke, J., Rice, S., Sacco, S., Squassante, L., Spiridigliozzi, G., Visootsak, J., Heller, J. & Khwaja, O. (2015) Assessment of Cognitive Scales to Examine Memory, Executive Function and Language in Individuals with Down Syndrome: Implications of a 6-month Observational Study. Front Behav Neurosci, 9, 300.

36. Pennington, B.F., Moon, J., Edgin, J., Stedron, J. & Nadel, L. (2003) The neuropsychology of Down syndrome: evidence for hippocampal dysfunction. Child Dev, 74, 75–93.

37. Zeleznikow-Johnston, A.M., Renoir, T., Churilov, L., Li, S., Burrows, E.L. & Hannan, A.J. (2018) Touchscreen testing reveals clinically relevant cognitive abnormalities in a mouse model of schizophrenia lacking metabotropic glutamate receptor 5. Sci Rep, 8, 16412.

38. Lim, J., Kim, E., Noh, H.J., Kang, S., Phillips, B.U., Kim, D.G., Bussey, T.J., Saksida, L., Heath, C.J. & Kim, C.H. (2019) Assessment of mGluR5 KO mice under conditions of low stress using a rodent touchscreen apparatus reveals impaired behavioural flexibility driven by perseverative responses. Mol Brain, 12, 37.

39. Itami, S. & Uno, H. (2002) Orbitofrontal cortex dysfunction in attention-deficit hyperactivity disorder revealed by reversal and extinction tasks. Neuroreport, 13, 2453–2457.

40. Oxelgren, U.W., Myrelid, A., Anneren, G., Ekstam, B., Goransson, C., Holmbom, A., Isaksson, A., Aberg, M., Gustafsson, J. & Fernell, E. (2017) Prevalence of autism and attention-deficit-hyperactivity disorder in Down syndrome: a population-based study. Dev Med Child Neurol, 59, 276–283.

41. Richards, C., Jones, C., Groves, L., Moss, J. & Oliver, C. (2015) Prevalence of autism spectrum disorder phenomenology in genetic disorders: a systematic review and meta-analysis. Lancet Psychiatry, 2, 909–916.

42. Warner, G., Moss, J., Smith, P. & Howlin, P. (2014) Autism characteristics and behavioural disturbances in ~ 500 children with Down’s syndrome in England and Wales. Autism Res, 7, 433–441.

43. Rowland, N.E. (2007) Food or fluid restriction in common laboratory animals: balancing welfare considerations with scientific inquiry. Comp Med, 57, 149–160.

44. Toth, L.A. & Gardiner, T.W. (2000) Food and water restriction protocols: physiological and behavioral considerations. Contemp Top Lab Anim Sci, 39, 9–17.

45. Goltstein, P.M., Reinert, S., Glas, A., Bonhoeffer, T. & Hubener, M. (2018) Food and water restriction lead to differential learning behaviors in a head-fixed two-choice visual discrimination task for mice. PLoS One, 13, e0204066.

46. Forestell, C.A., Schellinck, H.M., Boudreau, S.E. & LoLordo, V.M. (2001) Effect of food restriction on acquisition and expression of a conditioned odor discrimination in mice. Physiol Behav, 72, 559–566.

47. Tucci, V., Hardy, A. & Nolan, P.M. (2006) A comparison of physiological and behavioural parameters in C57BL/6J mice undergoing food or water restriction regimes. Behav Brain Res, 173, 22–29.

48. Guedj, F., Pennings, J.L., Ferres, M.A., Graham, L.C., Wick, H.C., Miczek, K.A., Slonim, D.K. & Bianchi, D.W. (2015) The fetal brain transcriptome and neonatal behavioral phenotype in the Ts1Cje mouse model of Down syndrome. Am J Med Genet A, 167A, 1993–2008.

